## Supplementary Figures (combined) for "Widespread perturbation of ETS factor binding sites in cancer"

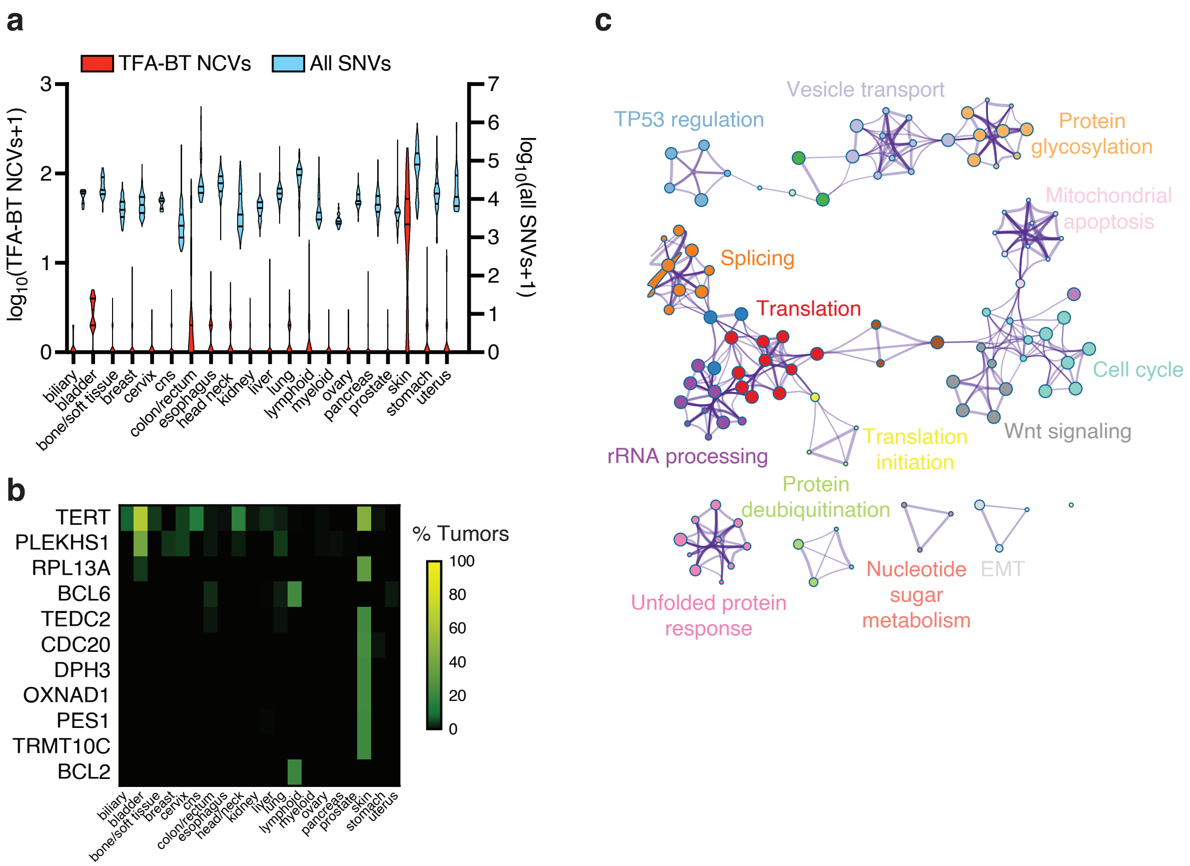


**Supplementary Figure 1. Frequency of TFA-BT NCVs across cancers.** (**a**) Number of TFA-BT NCVs and total SNVs per patient for each cancer type. (**b**) Percentage of tumors per cancer type with TFA-BT mutations for each of the indicated genes. Only genes with TFA-BT NCVs in at least 5% of tumor samples in at least one cancer type are shown. (**c**) Metascape enrichment network showing gene ontologies significantly associated with TFA-BT genes. Each node represents a gene ontology term. Pairs of nodes with Kappa similarities aboved 0.3 are connected by edges. and connected nodes share genes.

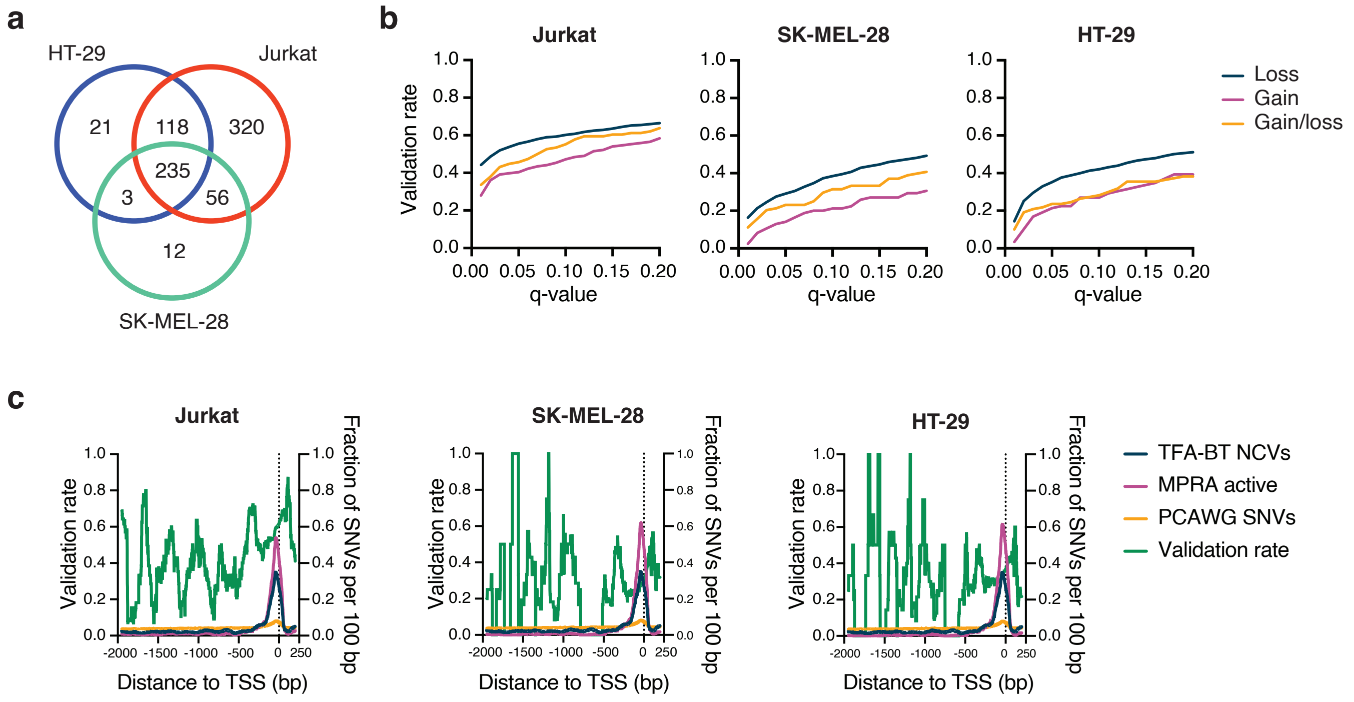


**Supplementary Figure 2. TFA-BT NCVs validation rate.** (**a**) Venn diagram showing the overlap between TFA-BT NCVs with significant allelic skew ( q < 0.05) by MPRAs identified in Jurkat, HT-29, and SK-MEL-28 cells. (**b**) Fraction of TFA-BT NCVs associated with loss, gain, or gain and loss of TFBSs within MPRA active regions that show expression allelic skew at different q-value thresholds in Jurkat, SK-MEL-28, and HT-29 cells. (**c**) MPRA validation rate of TF-ABT NCVs based on the genomic distance to transcription sart site (TSS). The fraction of NCVs per 100 bp for TFA-BT NCVs, MPRA active TF-ABT NCVs, and SNVs in the PCAWG cohort are also indicated.

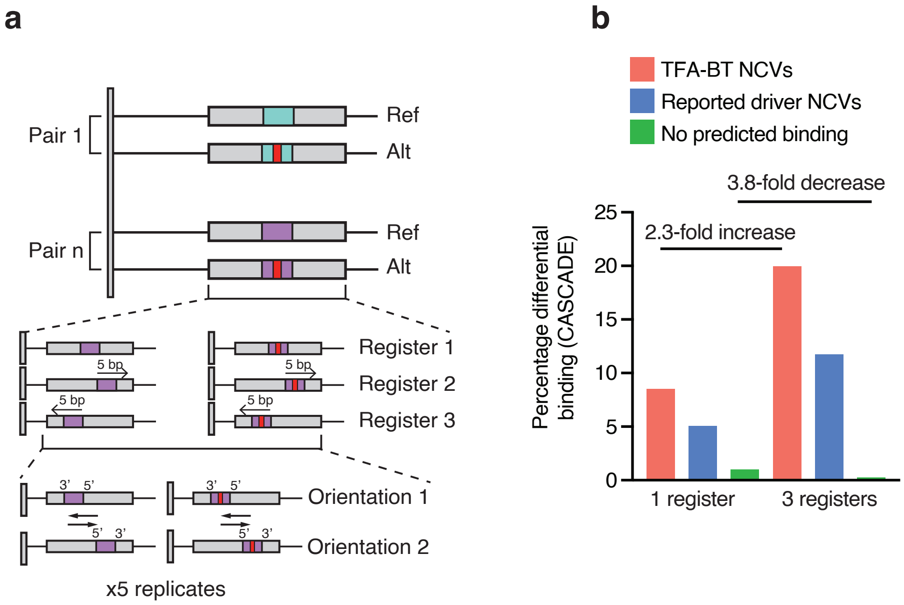


**Supplementary Figure 3. CASCADE experiment design**. (**a**) Outline of CASCADE experiments. Each reference (Ref) and alternative (Alt) NCV allele was tested in both orientations relative to the glass slide and using three registers: center, 5 bp shift upstream and 5 bp shift downstream. Each of these sequences was tested in five replicates placed in different positions in the array. (**b**) CASCADE performance improvement using three versus one register. The bar graph depicts the percentace of TFA-BT, reported driver, and no prodicted binding NCVs that show differential cofactor recruitment when tested in one or three registers.

Supplementary Figure 4

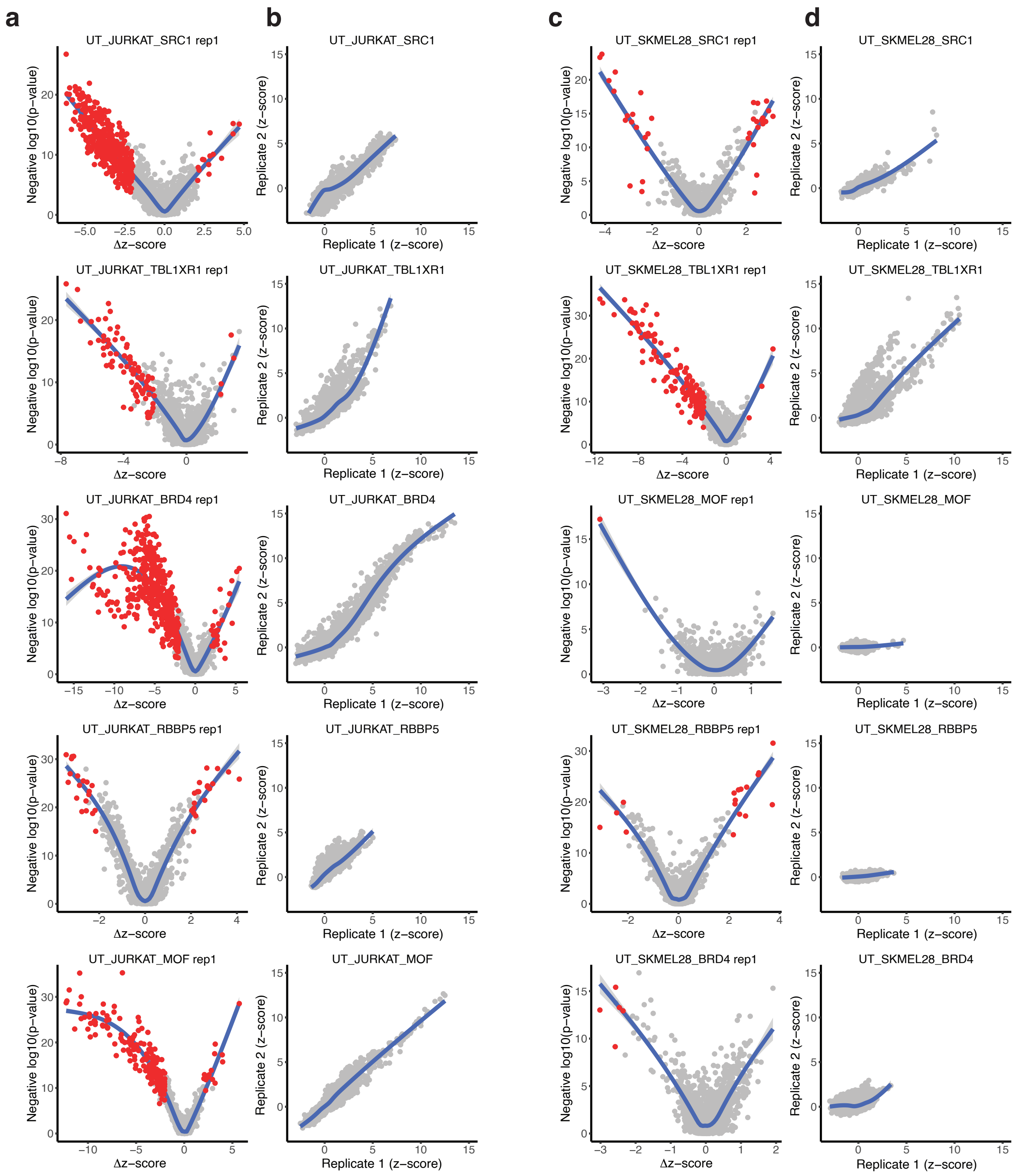


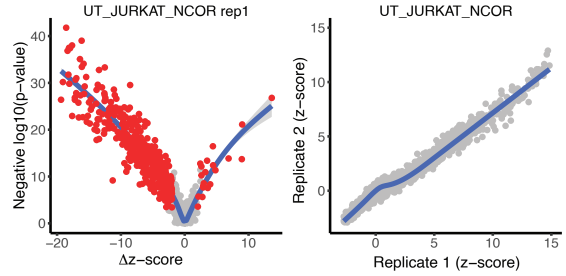


**Supplementary Figure 4. Identification of TFA-BT NCVs that alter COF recruitment.** (**a**, **c**) Impact of TFA-BT NCVs on COF recruitment using Jurkat (**a**) and SK-MEL-28 (**c**) nuclear extracts. Impact is quantified with -$\log_{10}$(p-value) of the COF recruitment to the different probe sets and the difference in PBM-determined Z-score between Ref and Alt alleles (Δz-score). The NCVs identified as significant are highlighted in red. (**b, d**) Reproducibity of CASCADE. Z-scores for each TFA-BT NCV are compared between biological replicates performed using Jurkat (**b**) and SK-MEL-28 (**d**) nuclear extracts.

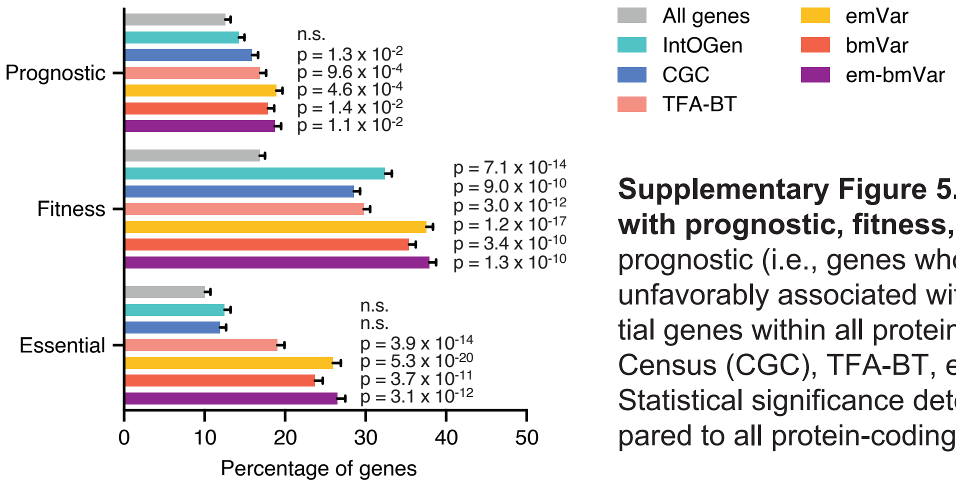


**Supplementary Figure 5. emVar and bmVar gene association with prognostic, fitness, and essential genes.** Percentage of prognostic (i.e., genes whose expression levels are favorably or unfavorably associated with cancer), fitness-related, and essential genes within all protein-coding, IntOGen, Cancer Gene Census (CGC), TFA-BT, emVar, bmVar, and em-bmVar genes. Statistical significance determined by Fisher's exact test compared to all protein-coding genes.

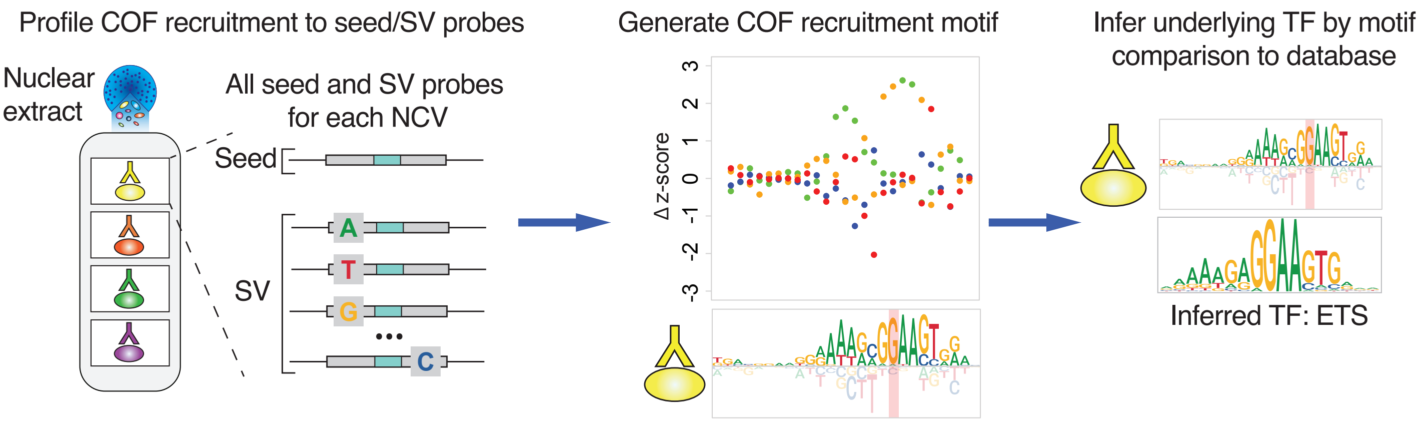


**Supplementary Figure 5. Identification of underlying DNA motifs affecting COF recruitment.** Outline illustrating the CASCADE approach to determine the DNA motifs affecting COF recruitment. COF recruitment is assayed to a “seed” probe containing the Ref or Alt NCV sequences in the genomic context and all single variant (SV) probes. The confetti plots show COF recruitment preferences to single variant probes along DNA sequence. Preferences are transformed to a COF recruitment motif. COF recruitment motifs are matched to TF motif databases to infer the identity of the TF recruiting the COF.

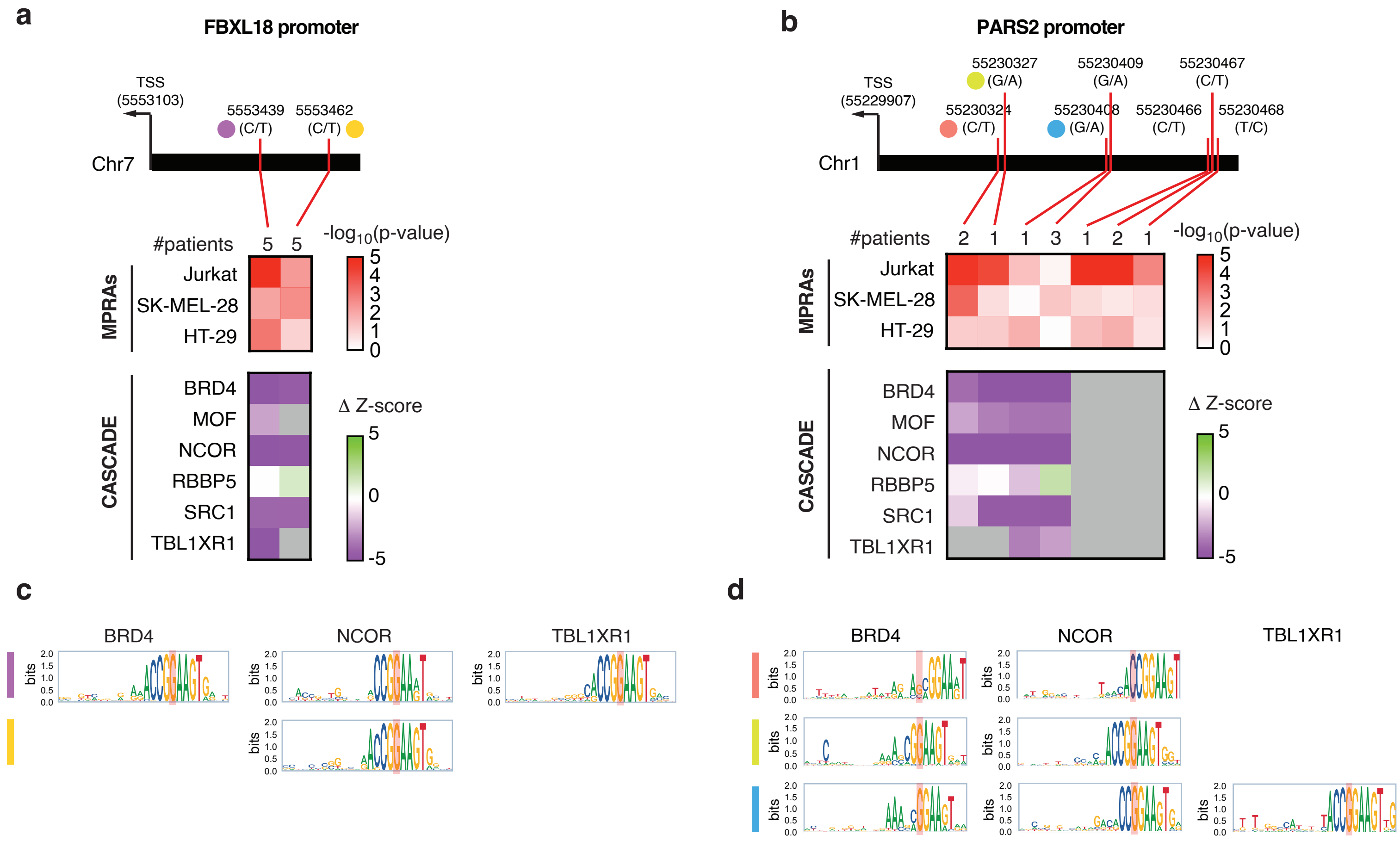


**Supplementary Figure 7. Altered transcriptional activity and COF recruitment within promoters.** (**a-b**) Changes in MPRA activity and COF recruitment for TF-ABT NCV in the (**a**) FBXL18 and (**b**) PARS2 promoters. The top heatmaps show the log10(p-value) of expression allelic skew in MPRA in Jurkat, SK-MEL-28, and HT-29 cells is indicated. The bottom heamaps show the altered COF recruitment by CASCADE, which is indicated as Δz-score. Gray cells indicate cases where the COF was not recruited to either NCV allele. Numbers at the top of the heatmaps indicate the number of patients in PCAWG carrying the indicated NCV. (**c-d**) COF recruitment motifs determined by single nucleotide variant scanning using CASCADE for the NCVs indicated in **a-b**.

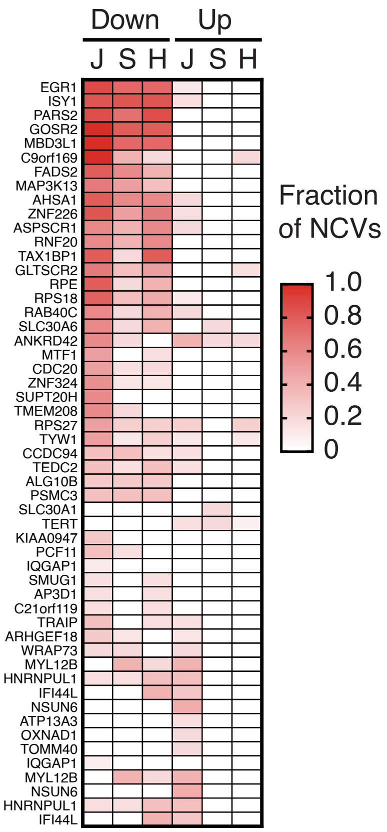


**Supplementary Figure 8. TFA-BT NCVs associated with transcriptional activation and repression.** Each cell in the heatmap represents the number of TFA-BT NCVs with increased (up) or decreased (down) transcriptional activity in MPRAs relative to the number of TFA-BT NCVs in MPRA active regions for the indicated gene and indicated cell line. J – Jurkat, S – SK-MEL-28, H – HT-29. Only genes with at least four TF-ABT NCVs in MPRA active regions are shown.
