## Supplementary Table 5 for "Widespread perturbation of ETS factor binding sites in cancer"

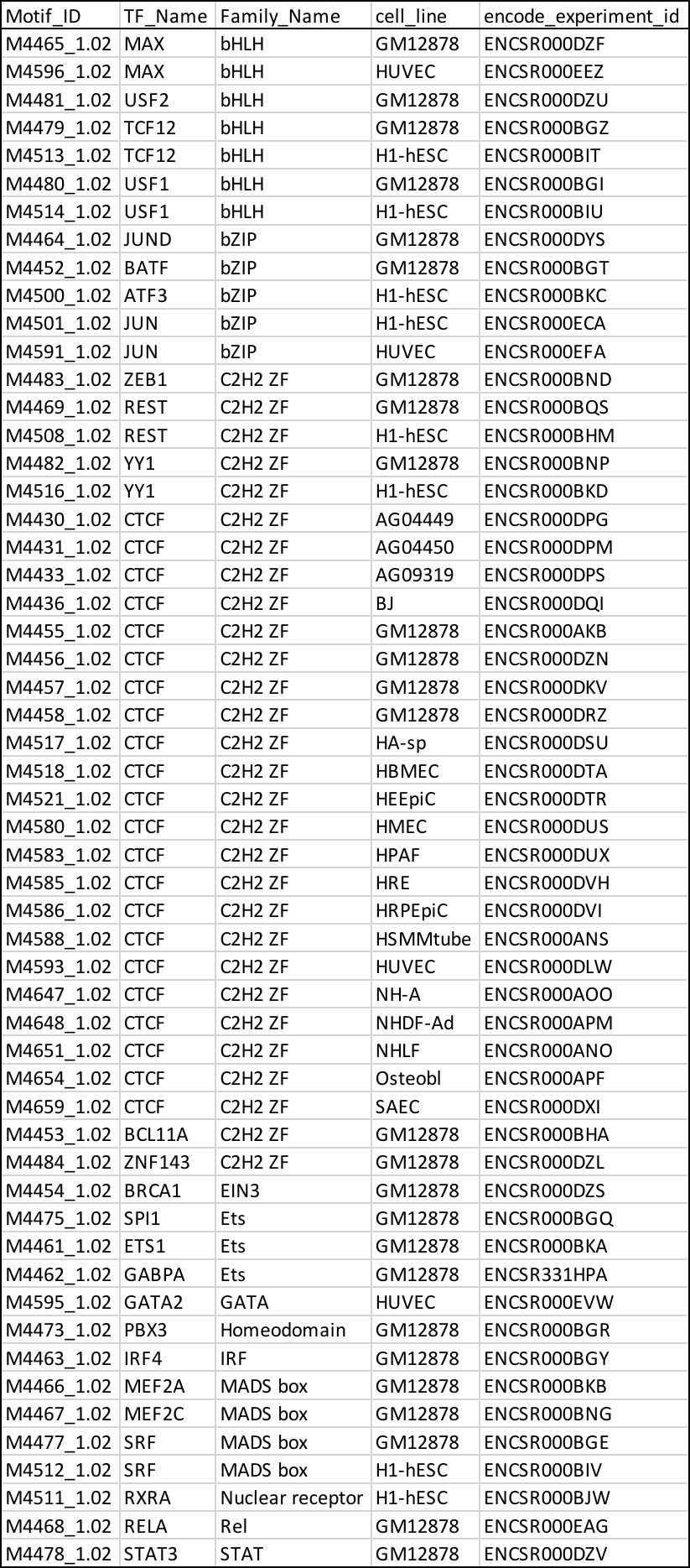

Supplementary Table 5: ChIP-seq experiments downloaded from ENCODE for allelic imbalance analysis. Motif ID – CIS-BP ID of motif used for predictions of differential binding; TF_name – name of TF associated with the indicated motif; Family_Name – TF family; cell_line – cell line corresponding to the ChIP-seq data from ENCODE; encode_experiment_id – ENCODE ID corresponding to the ChIP-seq experiment.
