## Supplementary Table 7 for "Widespread perturbation of ETS factor binding sites in cancer"

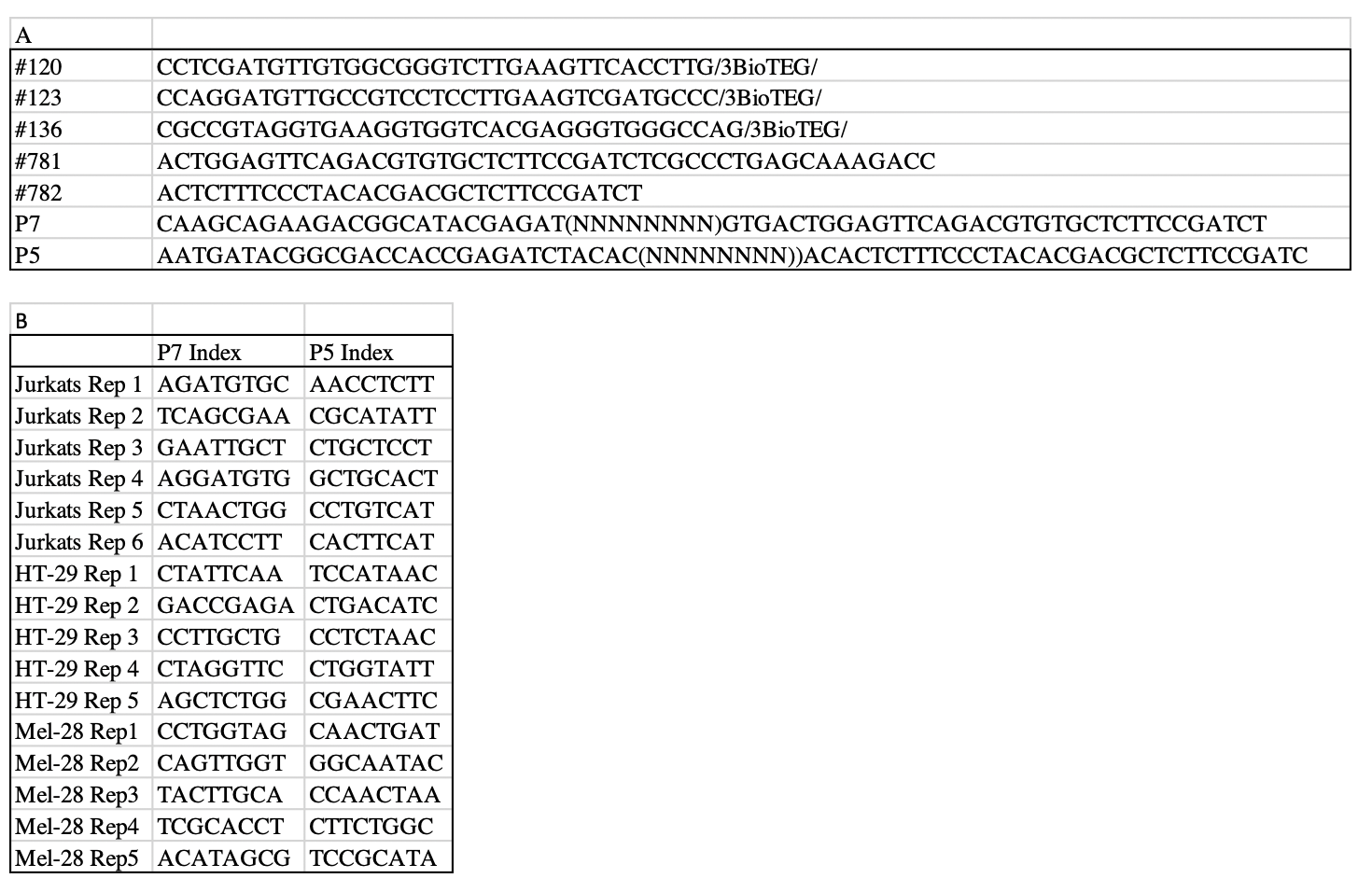

Supplementary Table 7: Primers and sequencing indices used. (A) Primers used in MPRA experiments and (B) Illumina Adaptor/Index sequences.
